# Screening bioactive food compounds in honey bees suggests curcumin blocks alcohol-induced damage to longevity and DNA methylation

**DOI:** 10.1101/2021.05.05.442791

**Authors:** Erik M. K. Rasmussen, Kristine L. Seier, Ingrid K. Pedersen, Claus Kreibich, Gro V. Amdam, Daniel Münch, John Arne Dahl

## Abstract

Various bioactive food compounds may confer health and longevity benefits, possibly through altering or preserving the human epigenome. While bioactive food compounds are widely being marketed as ‘improving health and longevity’ by counteracting harmful effects of poor nutrition and lifestyle, claimed effects are often not adequately documented. Using the honey bee (*Apis mellifera*) as a model species, we here employed a multi-step screening approach to investigate seven compounds for effects on lifespan and DNA methylation using ELISA and whole genome bisulfite sequencing (WGBS). A positive longevity effect was detected for valproic acid, isovaleric acid, and cyanocobalamin. For curcumin, we found that lifespan shortening caused by ethanol intake, was restored when curcumin and ethanol were co-administered. Furthermore, we identified region specific DNA methylation changes as a result of ethanol intake. Ethanol specific changes in DNA methylation were fully or partially blocked in honey bees receiving ethanol and curcumin together. Ethanol-affected and curcumin-blocked differentially methylated regions covered genes involved in fertility, temperature regulation and tubulin transport. Our results demonstrate fundamental negative effects of low dose ethanol consumption on lifespan and associated DNA methylation changes and present a proof-of-principle on how longevity and DNA methylation changes can be negated by the bioactive food component curcumin. Our findings provide a fundament for further studies of curcumin in mice and humans and offer an avenue to explore regarding possible prevention of health issues related to alcohol consumption.

## Introduction

Once thought to be a stable epigenetic mark, it is now widely accepted that DNA methylation can be dynamic. Recent advances in uncovering enzymatically oxidative derivates of 5-methylcytosine and a deeper understanding of the well-known, but poorly described demethylation of the paternal genome just after fertilization have led to a rising interest in the field of functional epigenetics and especially dynamic DNA methylation. The recognition of the dynamic nature of epigenetic marks provides a foundation for assessing the effect the environment and specific compounds have on the epigenome of living organisms.

Micronutrients and other bioactive food components hold potential as human epigenome effecting drugs (Anderson, Sant, & Dolinoy, 2012; McGowan, Meaney, & Szyf, 2008; Park, Friso, & Choi, 2012). The diversity of food supplements that is marketed for potential health benefits includes vitamins, plant extracts and synthetic analogues to natural compounds. While health effects are indisputable for many supplements, claimed benefits are unclear for others despite traditional beliefs. In addition, modes of actions are often elusive when concerning potential epigenetic effects. The assessment and understanding of epigenetics in metabolic studies is still in its infancy. Folic acid (vitamin B9) however, is an exceptional example of a compound with demonstrated inherent linkage to epigenomic mechanisms. Folic acid is an important precursor that can feed into the methyl donor pathway of DNA methylation (Anderson et al., 2012). It is already established as prophylactic drug during pregnancy to protect the fetus against neural tube defects (MRC VITAMIN STUDY RESEARCH GROUP 1, 1991). Sodium butyrate (mostly produced in the intestine from fermentation of dietary fiber, but also present in some dairy products) was one of the first bioactive food components shown to interfere with the posttranslational modifications on the N-terminal tails of core histones by acting as an inhibitor of certain histone modifying enzymes (Davie, 2003; Kruh, 1982; Vidali, Boffa, Bradbury, & Allfrey, 1978).

Although variations of the CRISPR/Cas9 system have been engineered to specifically modulate the methylome(Lei et al., 2017; Morita et al., 2016) (Liu et al., 2016; Vojta et al., 2016), these techniques are not readily available for human therapy as they would either need a very specific target vehicle, or require germline editing, which is a very controversial topic (Normile, 2018) (Lanphier, Urnov, Haecker, Werner, & Smolenski, 2015; The, 2018). Nevertheless, tissue or cell specific epigenetic editing may hold promises for future applications in human medicine, however, this is currently not achievable nor considered safe. Still, several bioactive food components, such as micronutrients and plant derived compounds, are deemed as safe for human consumption in relevant doses(Dashwood & Ho, 2007; Park et al., 2012). Some of these may confer health and longevity benefits, possibly through altering or preserving the human epigenome.

The synthetic compound valproic acid, an analogue of valeric acid derived from the plant *Valeriana officinalis* has for decades been used in treatment for epileptic seizures (for a review see (Ghodke-Puranik et al., 2013)). Hints about the mode of action was provided in the early 2000s when Phiel and coworkers (2001)(Phiel et al., 2001) discovered that valproic acid could inhibit histone deacetylases. Plant extracts are usually a mixture of different compounds, and improper storage may lead to formation of breakdown products like in the case of *Valeriana officinalis*, isovaleric acid(Eadie, 2004; Wills & Shohet, 2009). The effects of isovaleric acid on the epigenome remains largely unstudied.

A search for “curcumin”, the main compound isolated from the rhizome of the turmeric plant, on clinicaltrials.gov reveals over 200 clinical studies (as of 18.12.2020). Curcumin has been shown to confer some level of protection to the liver against damages from alcohol abuse in mice (Varatharajalu et al., 2016). However, further studies are needed. Curcumin has been implicated in histone hypoacetylation (Balasubramanyam et al., 2004; Kang, Chen, Shi, Jia, & Zhang, 2005). Considerable crosstalk between post-translational modifications of histones and DNA methylation has been demonstrated (Du, Johnson, Jacobsen, & Patel, 2015; Nan et al., 1998; Ooi et al., 2007).

However, low bioavailability of curcumin and usage of poor solvents (i.e. water) plague many of the earlier studies (Burgos-Moron, Calderon-Montano, Salvador, Robles, & Lopez-Lazaro, 2010; Nelson et al., 2017). These are challenges that must be considered when conducting in vivo studies.

Ethanol, one of the most harmful substances abused by humans in Europe, is also a versatile solvent of many organic compounds, including bioactive food components (van Amsterdam, Nutt, Phillips, & van den Brink, 2015). However, rodent model organisms like mice display large strain variations in ethanol tolerance and development of conditioned taste aversion (Broadbent, Muccino, & Cunningham, 2002; Crabbe et al., 2019; Rhodes et al., 2007; Risinger & Cunningham, 1995).

Scoring of survival in a model organism can be a powerful screening assay to directly determine advantageous or deleterious effects of plant-derived compounds fed at a range of concentrations. Promising hits can be further studied by whole genome bisulfite sequencing (WGBS) to assess the whole genome for DNA methylation changes associated with consumption of a compound. This strategy can uncover compounds that confer health and longevity benefits and provide insight into molecular mechanisms and pathways related to these benefits. Arguably, the honey bee is a cost effective and, due the short life span, an efficient model organism for such studies.

The honey bee is a demonstrated excellent model for functional epigenetics, specifically reversal of DNA methylation patterns (Erik Magne Koscielniak Rasmussen & Amdam, 2015). The honey bee hive is a highly complex structure with functionally sterile worker bees making up the bulk of the bees, communicating and performing specialized task to support the hive (Seeley, 1995). Importantly, unlike Dipterans like Drosophila melanogaster, honey bees have a functional DNA methylation system similar to humans (Erik Magne Koscielniak Rasmussen & Amdam, 2015; Wang et al., 2006). Herb and co-authors (2012)(Herb et al., 2012) have shown that a social perturbation of the DNA methylome in honey bees is possible, reversing previously established DNA methylation patterns when older worker bees revert to tasks usually performed by younger workers.

In addition to its semi-domesticated hive habitat, honey bees can be housed inside the laboratory with relatively little effort or use of lab space (Huang et al., 2014). This allows tight control of food intake, and usage of many individuals facilitates screening of different compounds at a range of concentrations. The honey bee diet is also exclusively plant derived, consisting mainly of nectar and pollen. This contrasts with rodent model organisms which in the wild are opportunistic eaters and can consume smaller insects and easily consumable meat resources. In addition, honey bees could potentially encounter ethanol in floral nectars, as yeast have been found in nectar that bees forage (de Vega, Herrera, & Johnson, 2009).

In this study we use a three-step screening approach were seven compounds, at a range of concentrations, were first screened for effects on longevity. All compounds and concentrations were assessed for any deviation in consumption, and only compounds that did not affect food consumption were included in further analysis. These compounds were then screened for global methylation changes. The most promising compound was selected for WGBS and in-depth analyses of how DNA methylation may be affected by substance exposure. A positive survival effect was detected for valproic acid, isovaleric acid, and cyanocobalamin. We show that ethanol has a detrimental effect on worker bee lifespan and induces changes in DNA methylation patterns. For curcumin, we found that lifespan shortening caused by ethanol intake, was restored when curcumin and ethanol were co-administered. Furthermore, we identified region specific DNA methylation changes as a result of ethanol intake. Ethanol specific changes in DNA methylation were fully or partially blocked in honey bees receiving ethanol and curcumin together. We present a proof-of-principle on curcumin restoring alcohol-associated shortened lifespan and DNA methylation changes in an important model organism. Our findings provide a fundament for further studies of curcumin in mice and humans and offer an avenue to explore regarding possible prevention of health issues related to alcohol consumption.

## Material and methods

### Animals

Honey bees (*Apis mellifera carnica* Pollmann) were kept at the Norwegian University of Life Sciences’ apiary at Aas, Norway. Honey bee studies on survival and feed consumption were carried out during fall of 2013, specimen for DNA methylation studies were collected during winter 2014 (see below).

To establish similar life history traits among all tested specimens we exclusively used one worker type: winter bees. To achieve this, we used established tools to manipulate the hive’s social demography and worker bee tasks (Daniel Münch et al., 2013). In brief, the characteristic feature of winter (*diutinus*) bee colonies is the absence of brood, which triggers physiological changes in workers, e.g., by diverting resources into increased nutrient storage, winter worker types typically show higher resilience and longer survival (Fluri, 1990; Fluri, Lüscher, Wille, & Gerig, 1982) (Münch, Kreibich, & Amdam, 2013; Smedal, Brynem, Kreibich, & Amdam, 2009). To assure collection of winter worker type, all colonies for sampling in January were confirmed to be broodless prior to collection. To obtain broodless hives and winter worker types already in fall (September-October), we removed all brood combs with eggs and larvae at least 2 weeks prior collection, and queens were prevented from egg-laying through confining them in queen cages(D. Münch et al., 2013).

### Survival assays and assessment of feed consumption

Sample collection in fall, was essentially carried out as follows: honey bees were collected from two individual donor colonies (‘colony replicate’) in two rounds, each separated by two to three days (‘day replicate’). From each donor colony, four cages with approximately 50 bees in each cage were collected, resulting in a total of 16 cages and approximately 800 bees for each round.

For the entire subsequent test period, bees remained in the plastic cages, which had the following dimensions: width/depth/height: 19×18×23 cm and were lined with patches of mesh for ventilation. After collection, cages were transferred into a climate chamber, set to temperature of 26°C with approximately 60% relative humidity (e.g. (Dickel, Munch, Amdam, Mappes, & Freitak, 2018) (Huang et al., 2014). For acclimation and to control for possible effects of sample collection on initial mortality rates, all cages received the control diet (see below) for the first two days of captivity. Bees that died during this initial period were removed and not considered in survival statistics.

For subsequent survival and food consumption testing bees had ad libitum access to a water feeder and another feeder containing the respective treatment diet. Treatment identity (factor: dietary supplementation) was balanced with respect to day and colony replicates, and was otherwise randomly assigned to the individual cages. Each day, dead bees were removed and the number of dead individuals per cage was recorded. Similarly, food and water feeders were changed, and the consumed volume was recorded daily. After 41 days the experiment was terminated, and any remaining bees were censored.

The average daily consumption per bee for the first 10 treatment days was calculated per cage in order to investigate if any lifespan differences could be attributed to differences in consumption.

We have tested for possible effects of dietary supplementation with isovaleric and valproic acid, sodium butyrate, cyanocobalamin, folic acid, ethanol, and curcumin. All treatment diets were made based on the same stock feed solution, which also served as the control diet. The stock feed solution contained 50 % Bifor® (Nordic Sugar; 37 % sucrose, 19 % glucose, 19 % fructose in water), 2% Grace’s amino acid mix for insect cell culture (Sigma-Aldrich cat no G0148) or RPMI 1640 (Sigma-Aldrich cat no R7131), 1% lipid mix for insect cell culture (Sigma-Aldrich cat no L5146) and 47% dH_2_O. In treatment diets the dH_2_O fraction of the stock feed was partly substituted by the respective compound (details in Supplementary Table 1). All tested compounds except ethanol were given at 3 concentrations (considered low, medium, high concentration based on literature) with a 10x concentration increment. Each treatment (factor: concentration) was given to four replicate cages with 50 bees each, making a total of 3*4=12 cages plus 16 (for isovaleric acid, valproic acid, folic acid, and cyanocobalamin) or four (for sodium butyrate, ethanol, and curcumin) control fed cages (50 bees each) per tested compound.

### Preparation of bees for molecular testing

Honey bees were collected from two separate donor hives on January 15^th^, 2014. In total, 18 cages with approximately 50 bees each were collected from both hives resulting in approximately 900 bees.

All honey bees were fed a control diet with the same constituents as the survival assay experiment for the first 2 days of captivity. After 2 days, food diets were exchanged with the same treatment diets used in the survival assay experiment. For a detailed makeup of diets used see Supplementary table 1. Each treatment diet was given to two cages to reduce the chance of bias due to cage effects.

Temperature, humidity, number of dead honey bees, food and water consumption was recorded daily. Fresh food and water were also given daily along with removal of any dead bees. After 10 days of treatment diet feed, all remining honey bees were flash frozen in liquid nitrogen and stored at – 80 ° C until further processing.

### DNA extraction

DNA was extracted essentially as described previously (Erik M. K. Rasmussen et al., 2016). The stinger along with the gut was pulled out of the separated abdomen and discarded. The remaining abdominal carcass with the fat body tissue was used for further analyses. Briefly, abdomen carcasses with adhering fat body were homogenized in ATL buffer (QIAGEN 19076) containing 0.4 mg Proteinase K (Sigma Aldrich P8044). Samples were then incubated at 56 °C with shaking at 400 rpm for 16 h in a Thermomixer (Eppendorf 5355 000.011). The lysate (500 µl) was extracted once in phenol:chloroform:isoamylalcohol (25:24:1) before the aqueous phase was RNAse treated by incubating the sample with 5 µl of RNAse A (20 mg/ml) at 37 °C with shaking at 550 rpm for 30 min. The aqueous phase was extracted again using phenol:chloroform:isoamylalcohol (25:24:1) before genomic DNA was precipitated by adding 1/10 volume equivalents of 3 M sodium acetate (pH 5.3), 10 µl of linear acrylamide (ThermoFisher Scientific AM9520) and 2.5 vol of ethanol, washed twice in 70 % ethanol, air dried, and dissolved in nuclease free water.

### Assessment of global methylation state using 5mC specific ELISA

Enzyme linked (ELISA) was performed using the Zymo Research 5-mC DNA ELISA kit according to the manufacturer’s instructions. Briefly, 100 ng of DNA was diluted with 5-mC coating buffer to a volume of 100 µl. The DNA was denatured at 98 °C for 5 min and immediately incubated on ice for 10 min. Denatured DNA was transferred to wells and incubated in darkness at 37 °C for 1 h, allowing the DNA to settle and adhere to the wells. After incubation, the buffer was discarded, the plate was washed three times with 5-mC ELISA buffer, and kept with the 5-mC ELISA buffer at 37 °C for 30 min in darkness. After incubation, the buffer was discarded, and primary and secondary Horseradish Peroxidase conjugated antibodies were added to each well for labeling of 5mC (1:2000 and 1:1000 concentrations respectively). The plate was incubated at 37 °C for 1 h in darkness. After incubation, the plate was washed three times with 5-mC ELISA buffer, HRP-developer was added and allowed to develop for 45 min before absorbance at 405 nm was measured with a microplate reader. The percentage of 5mC present in the samples were calculated according to the manufacturer’s description.

### Whole genome bisulfite sequencing library preparation

Sequencing libraries were constructed essentially as described before (Urich, Nery, Lister, Schmitz, & Ecker, 2015). Briefly, 2 µg of genomic DNA spiked with 10 ng of λ-phage DNA which is devoid of DNA methylation was fragmented using a Sartorius Labsonic M sonicator, with cycle settings at 0.5 and power/amplitude set to 30 %. Average fragment length was assessed on an Agilent Bioanalyzer DNA High Sense chip. Lower and upper size cut-offs were performed using Agencourt AMPure XP beads at a volume of 0.6x and 1.4x to DNA solution respectively. Sticky ends were repaired using Epicentre End-it DNA end repair kit according to the manufacturer’s recommendations. Poly-Adenylated tails were added to blunt end fragments using New England Biolabs (NEB) Klenow Fragment (3’-->5’ exo-) and NEBNext® dA-Tailing Reaction Buffer according to the manufacturer’s recommendations. The reactions were cleaned up using Agencourt AMPure XP beads at a volume of 1.4x to DNA solution. Methylated adapters were ligated to eluted DNA using NEB T4 DNA ligase according to manufacturer’s recommendations. Ligation reactions were purified twice using Agencourt AMPure XP beads at a volume of 1.0x to DNA solution. Eluted DNA was bisulfite converted using Human Genetic Signatures MethylEasy™ Xceed kit according to the manufacturer’s recommendations. A low cycle PCR was performed to increase sequencing depth. To estimate the number of cycles needed, PCR reactions were quantified using KAPA Library Quantification Kit using the manufacturer’s recommendations. Completed PCR products were cleaned up using Agencourt AMPure XP beads at a volume of 1.0x to DNA solution. Libraries were pooled at equimolar ratios and sequenced on the HiSeq 2500 (Illumina).

### Alignment and identification of differentially methylated regions

Preliminary quality control was performed using FastQC (available from https://www.bioinformatics.babraham.ac.uk/projects/fastqc/). Low quality bases were removed using Trimmomatic (Bolger, Lohse, & Usadel, 2014). The bisulfite conversion efficiency was calculated by mapping the samples to the lambda phage genome. We set the conversion rate cut-off at 98 %. Trimmed reads were aligned to the Amel 4.5 reference genome (Elsik et al., 2014) using Bismark (Krueger & Andrews, 2011) and Bowtie 1(Langmead, Trapnell, Pop, & Salzberg, 2009). Mapped reads were then deduplicated to remove PCR bias. Cytosine report files where then imported into R (version 3.5.3) using bsseq (Hansen, Langmead, & Irizarry, 2012) (R Core Team, 2019). The identification of differentially methylated regions was performed essentially as before (Herb, Shook, Fields, & Robinson, 2018). Briefly, only cytosines with an average coverage of >= 6 in all samples and >= 10 % methylation levels in three or more samples were defined as methylated. Percentage measurements derived from ratios of reads containing methylated and unmethylated cytosines in the CpG context where then arcsine transformed. Suggestive CpGs were identified using Student’s t-test. Correction for multiple testing was performed using the qvalue package (available from https://github.com/StoreyLab/qvalue). Regions of individually methylated CpGs were designated by the bumphunter package (Jaffe et al., 2012). The raw p-values for each methylated CpG in each region were combined using the comb-p software package and corrected for multiple testing (Pedersen, Schwartz, Yang, & Kechris, 2012).

### Data analyses of survival and consumption assays

Survival data was analyzed, and plots were generated using the statistical software packages survival and survminer in R (version 3.5.3)[(Therneau & Grambsch, 2000), available from https://cran.r-project.org/package=survminer,(R Core Team, 2019). To assess overall effects among multiple treatment groups we used the log-rank test for trends. For pairwise comparison between treatment concentration and respective control we calculated log-rank statistics.

Corrections for multiple testing were done using the False Discovery Rate/Benjamini Hochberg Approach (Benjamini & Hochberg, 1995). Consumption data was analyzed using the STATISTICA 13 software package (Statistica, Dell Inc, Tulsa, USA). We used one-way ANOVA analysis to assess overall effects among multiple treatment groups. We used the Fisher LSD as a post-hoc test to determine which concentration(s) differed from the control(s) if any.

## Results and Discussion

### Screening of food bioactive compounds reveals effects on survival in honey bees

We selected seven food bioactive compounds for our study that are relevant regarding human exposure through food, dietary supplements, pharmaceuticals and abuse. We first asked if intake of the seven chosen compounds could affect the lifespan of honey bees. A three-step screening approach was applied, were 7 substances, at a range of concentrations, were first screened for effects on longevity. All substances and concentrations were assessed for any deviation in consumption, and only substances that did not affect food consumption were included in further analysis. These substances were then screened by ELISA for global methylation changes. The most promising candidate was selected for WGBS and in-depth analyses of how DNA methylation may be affected by substance exposure. For an overview of the flow of screening methodologies see figure 1. Honey bees are short lived and easy to keep in the laboratory. Henceforth, this enabled us to use just under 200 bees per tested compound concentration, a number of individuals that would be out of reach for most labs using rodent model organisms. Kaplan-Meier survival analyses on overall data (Log-rank test for trends) revealed significant treatment effects on survival for isovaleric acid (N _control/0.10mg/ml/1.0mg/ml/10.0mg/ml_ = 795/188/197/182; p<0.005), valproic acid (N _control/ 0.1 mg/ml/ 1.0 mg/ml/ 10.0 mg/ml_ = 795/194/192/193; p<0.005), cyanocobalamin (Vitamer of B12)(N _control/ 0.02 µg/ml/ 0.2 µg/ml/ 2µg/ml_ = 795/182/185/193; p<0.005) and folic acid (N _control/ 5 µg/ml/ 50 µg/ml/ 500 µg/ml_ = 795/192/193/191; p<0.005), but not for sodium butyrate (N _control/0.01mg/ml/0.1mg/ml/1.0mg/ml_ = 197/191/122/208; p>0.005). Pair-wise log-rank tests against controls with corrections for multiple testing were done to identify survival-effective concentrations for each compound (for an overview of all post-hoc tests see Figure 2A). We report longer survival for honey bees fed isovaleric acid at concentrations of 0.1 and 1 mg/ml (N _control/0.10mg/ml/1.0mg/ml/_ = 795/188/197; p_0.1mg/ml/1mg/ml_ <0.05), however, at 10 mg/ml we found survival to be shorter (N _control/10.0mg/ml_ = 795/182; p_10mg/ml_ <0.05)(Supplemental Figure 1). For valproic acid fed honey bees we observed longer survival for 0.1 mg/ml (N _control/0.1mg/ml_ = 795/194 p<0.05, Supplemental Figure 2), and found significantly shortened survival for higher concentrations (N _control/1.0mg/ml/10.0mg/ml_ = 795/192/193; p _1.0mg/ml/10.0mg/ml_ <0.05). Valproic acid is a potent anticonvulsant drug used in the treatment of multiple neurological and psychological diseases in humans (Ghodke-Puranik et al., 2013). However, its use often needs to be carefully monitored as numerous side-effects can occur. Our results indicate that high sensitivity for valproic acid is also conserved in honey bees. Cyanocobalamin given at a concentration of 0.2 µg/ml significantly shortened survival (N _control/0.2 µg/ml_ = 795/185, p<0.05, Supplemental figure 3). For folic acid (Vitamin B9) fed honey bees an effect on survival, i.e. shortened survival, was only found for the lowest concentration (5 µg/ml)(N _control/5 µg/ml_ = 202/2192; Supplemental figure 4). Folic acid and cyanocobalamin have received little attention in insects (S. A. Blatch, Stabler, & Harrison, 2015). However, there are indirect evidence that insects can obtain folate from gut bacteria (S. Blatch, Meyer, & Harrison, 2010). One may envisage that the gut microbiome might be shifted in a harmful way by exposure to folic acid rich diets, possibly leading to reduced fitness and survival.

**Figure 1:**
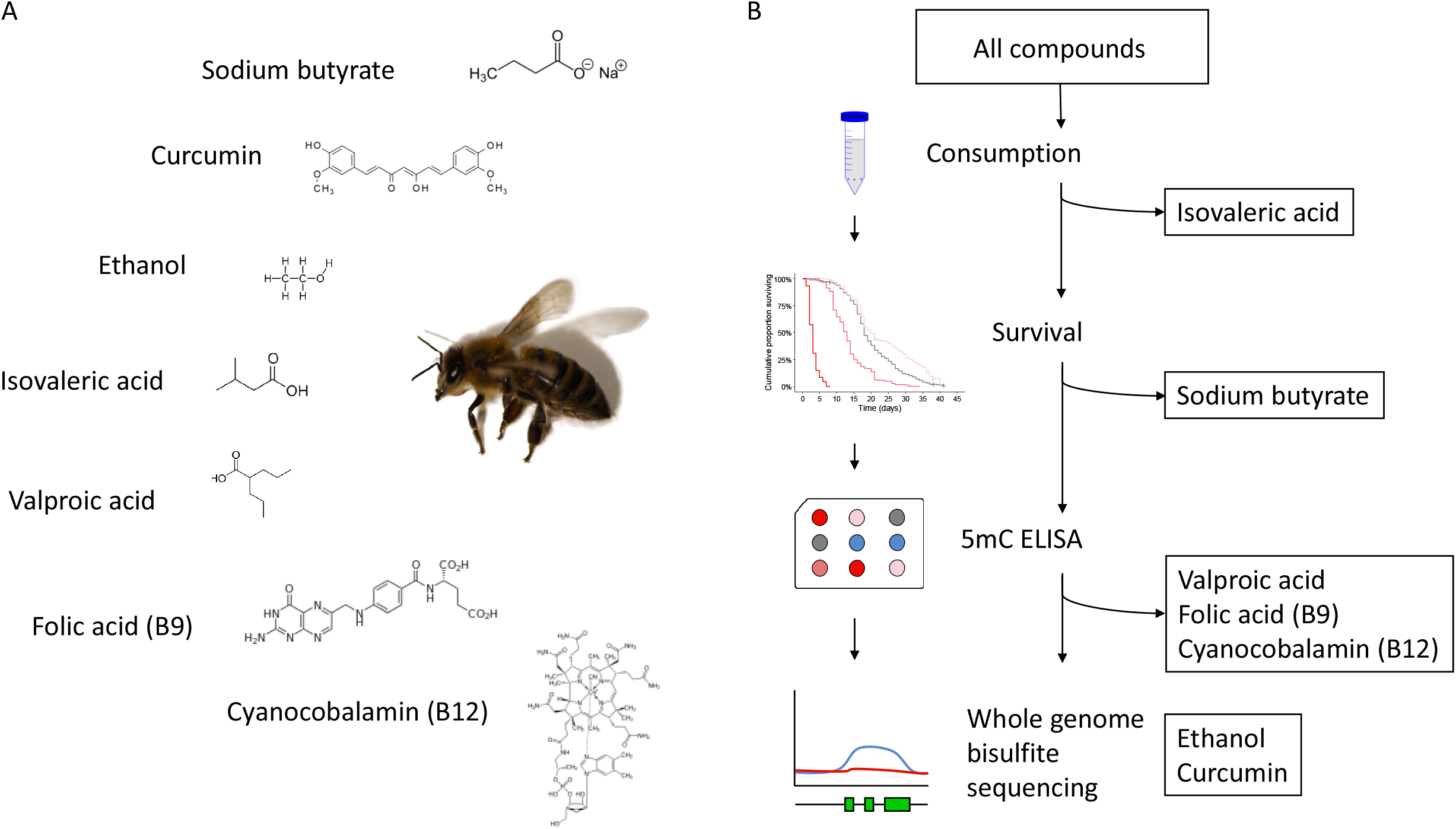
Overview of experimental setup. (A) Overview of the 7 initial compounds used. (B) Overview of flow of screening methodologies. A three-step screening approach was applied, were the 7 substances, at a range of concentrations, were first screened for effects on longevity. All substances and concentrations were assessed for any deviation in consumption, and only substances that did not affect food consumption were included in further analysis. These substances were then screened by ELISA for global methylation changes. The most promising candidate was selected for WGBS and in-depth analyses of how DNA methylation may be affected by substance exposure.

**Figure 2:**
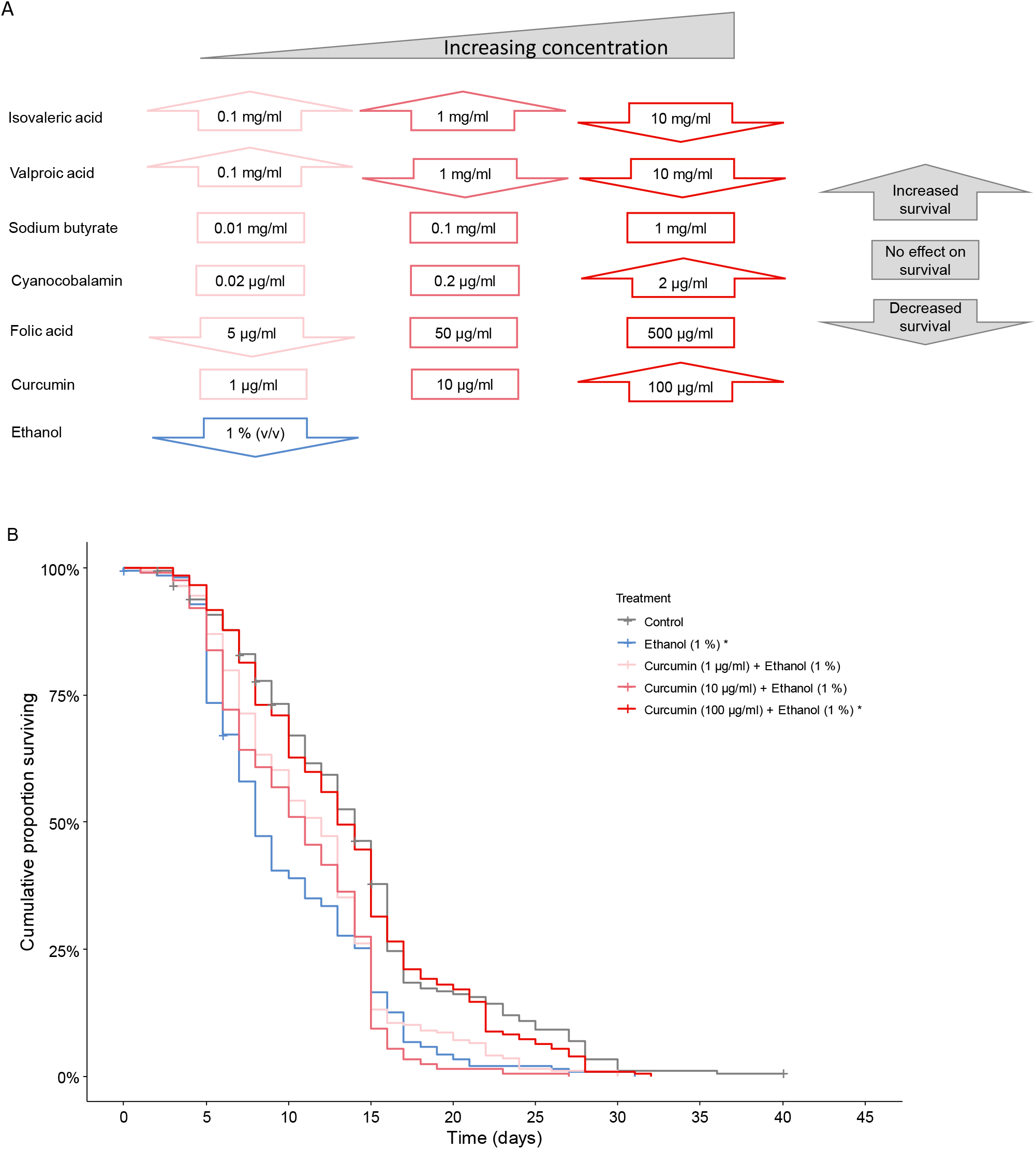
Effects of bioactive food compounds on honey bee survival. (A) Overview of all substances studied and the selected concentrations used in the feed. Upward pointing arrow signifies increased life span, rectangular box, no statistical effect on life span, downward pointing arrow signifies shorter life span (P < 0.05 Pair-wise Log-Rank test corrected using FDR). (B) Kaplan-Meier plot of survival curves of ethanol (1 % (v/v)) alone and in combination with curcumin at 1, 10 and 100 µg/ml. Pluss signs indicate censored honey bees. Asterisks in legends indicates significant effects on survival compared to controls.

In accordance with what has previously been reported, we show detrimental effects of ethanol on honey bee survival (N _control/ethanol_ = 198/208; p < 0.05, Figure 2B). Previous work reported a negative effect of ethanol on life span in honey bees for concentrations down to 2.5 % (v/v) (Mustard, Oquita, Garza, & Stoker, 2019). Using a longer-lived phenotype than Mustard et al. we observed reduced life span caused by an ethanol concentration as low as 1 % (v/v), indicating that low concentration of ethanol is more harmful than previously assumed.

Remarkably though, when honey bees were fed a diet that contained a high concentration of curcumin (100 µg/ml) together with ethanol (1 %, v/v), the life span was significantly increased as compared to ethanol intake only (Figure 2B)(N _ethanol / ethanol & curcumin 100 µg/ml_ = 208/204; p < 0.05). In fact, with 100 µg/ml curcumin in the ethanol containing diet, the ethanol associated reduction in life span was largely restored to that observed in the control (Figure 2B). However, honey bees fed with lower concentrations of curcumin (1 µg and 10 µg/ml) together with ethanol (1 % (v/v)) did not show a significant increase in life span (N _ethanol /10 µg/ml/1 µg/ml_ = 208/204/201; p > 0.05). This demonstrates the need for a relatively high concentration of curcumin for the favorable effect of restoring lifespan shortening caused by ethanol intake. Curcumin has previously been reported to increase lifespan in a study of infections in honey bees (Strachecka, Olszewski, & Paleolog, 2015). Taken together with our results, one may speculate that curcumin could possibly act through a common mechanism, for example by increasing stress resilience in honey bees.

In a survival study based on modifications of the feed composition one need to consider that differences in survival could potentially be affected by food palatability or other more adverse effects on food intake. To control for this, we monitored food consumption. Significant food consumption differences were only found for isovaleric acid (One-way ANOVA, N _control / 10 mg/ml / 1,0 mg/ml / 0,1 mg/ml_=4/4/4/4; F=29.76; p<0.001). A post-hoc test revealed that bees given the 10 mg/ml and 1 mg/ml diet consumed significantly less food (Fisher-LSD, p_1mg/ml_<0.05, p_10mg/ml_<0.001). Therefore, we cannot rule out that the observed differences in survival for feed containing 10 mg/ml and 1 mg/ml isovaleric acid was due to lower food intake. There was however no effect on food consumption for feed containing 0.1 mg/ml isovaleric acid. We observed no effect on food consumption for any of the other studied food bioactive compounds, and the observed effects on life span can therefore likely be attributed to the intake of the studied compounds. As we had to change the amino acid source during the experiment due to supplier constraints, we also investigated if the source of amino acids had any effect on survival. Differences in survival were indeed observed (Log-rank test, N _Grace’s aa mix / RPMI 1640_ = 198/795, p<0.0001) (Supplemental figure 5). However, we have not made any direct comparisons among compounds that use a different amino acid source for feed stock solutions.

### Assessment of effects of food bioactive compounds on global DNA methylation

Next, we asked if differences in lifespan after treatment with the tested compounds could be linked to changes in global DNA methylation. In order to reduce complexity, only two concentrations from each compound (along with their respective controls) were selected for assessment by ELISA (Supplementary Table1). Despite the relatively sparse methylated genome of the honey bee, ELISA based methods have successfully been applied to detect changes in global DNA methylation (Biergans, Giovanni Galizia, Reinhard, & Claudianos, 2015). For our assessed conditions we did not observe any significant effect on the global DNA methylation level among any of the tested compounds (One-tailed Mann-Whitney test; p>0.05). However, we did observe a trend towards hypermethylation when bees were fed ethanol alone. Notably, this trend was reversed in honey bees that had consumed the feed containing both ethanol (1 % (v/v)) and the highest curcumin concentration (100 µg/ml) (Supplementary figure S6)(One-tailed Mann-Whitney test; p=0.07446; n_1_= 4, n_2_=4). Locus-specific changes in DNA methylation may still occur although they cannot be identified when analyzing DNA methylation changes on a global level with the ELISA methodology that only has modest sensitivity.

### Exons are enriched for CpG methylation in honey bees

Encouraged by the observed trends in global DNA methylation changes in honey bees fed ethanol alone and co-fed ethanol and the highest curcumin concentration, we sat out to perform whole genome bisulfite sequencing (WGBS) in order to assess region specific DNA methylation changes throughout the honey bee genome. We carried out WGBS on DNA from abdominal extracts, which contain the fat body organ that is in part analogous to mammalian liver and white adipose tissue, thus important in energy metabolism, from a total of 13 honey bees, where five were fed 1 % (v/v) ethanol (n _ethanol_ = 5), three with 1% (v/v) ethanol combined with 100 µg/ml curcumin (n _curcumin_ = 3), and five controls (n _control_ = 5). Too low bisulfite conversion rates hindered the inclusion of two more of the curcumin and ethanol co-fed individuals for DNA methylation analysis. Assessment of bisulfite conversion rates and additional quality controls were carried out for all samples (Section M&M and Supplementary table 2).

As an added quality measure, we investigated if exons would be enriched for methylated CpGs, as methylated CpGs have been reported to be confined to exons in most insect species (For review see (Erik Magne Koscielniak Rasmussen & Amdam, 2015)). A CpG dinucleotide was included for analysis if the average coverage was six times or more, and that at least three samples showed a methylation level of 10 % or higher for this CpG. As expected, exons were enriched for methylated CpGs compared to whole mRNA sequence bins (Supplemental Figure 7). We then asked if any DNA methylation differences between treatments could be observed on a global scale using nucleotide resolution data. However, and as observed with the ELISA based method, no statistically significant differences could be found on the global level (One-tailed Mann-Whitney tests, n_control/ethanol/curcumin_ = 5/5/3 p > 0.05, Supplemental figure 8).

### Curcumin blocks alcohol associated DNA methylation changes

Using a previously established pipeline for honey bee methylomics (Herb et al., 2018), we identified seven hits for genomic regions that were significantly differentially methylated (Figure 4, Supplemental figures 9-11 and Supplemental table 2). Four regions were differentially methylated in ethanol exposed honey bees as compared to controls. Of these, three regions were hypermethylated and one hypomethylated. Three regions were differentially methylated (all hypomethylated) in curcumin and ethanol fed honey bees as compared to ethanol-only. (Supplementary Table 2). Upon further investigation, it became clear that three of the differentially methylated regions in curcumin and ethanol co-fed honey bees versus ethanol-only, overlapped with three of the differentially methylated regions observed in bees fed ethanol only. Ethanol induced hypermethylation in these three regions (Figure 3A-C), whereas curcumin remarkably protected against ethanol-induced hypermethylation in all of these regions.

**Figure 3:**
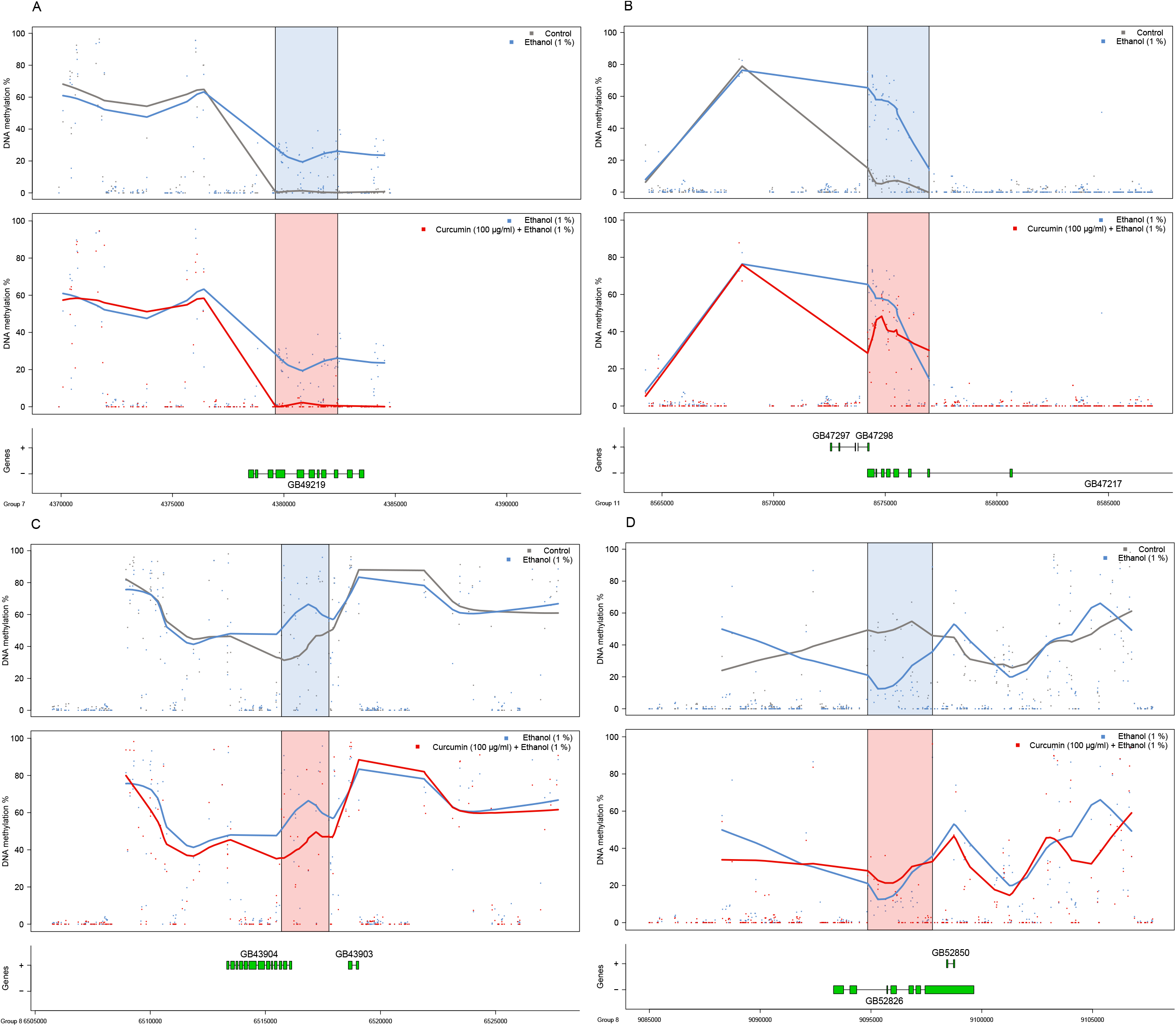
**(A-D)** Differentially methylated regions where the top panel depicts the situation in control (grey) versus ethanol (blue) fed honey bees, and the blue shaded area indicates the significant DMR. 10 kbp upstream and downstream (if no gaps exist in the Amel 4.5 genome build) are plotted for reference (white background). The middle panel depicts the comparison between ethanol-only (blue) and the honey bee group fed ethanol and curcumin (red). The lower panels depict the gene structure by exons (boxes) and introns (lines) by strand according to linkage groups.

Curcumin administered together with ethanol resulted in DNA methylation levels similar to the levels observed in the control honey bees for three out of four ethanol affected regions (Figure 3A-C and supplemental figures 9-11). These three ethanol-affected and curcumin-blocked differentially methylated regions are located in the gene bodies of the loci GB49219, GB47217 and GB43904, within linkage groups 7, 11 and 8, respectively. The Drosophila homologue of GB49219 is known as Gudu, which is necessary for male fertility (Cheng, Ip, & Xu, 2013). Based on this it would be interesting to investigate if male bees (drones) would experience transiently reduced sperm quality after excessive alcohol consumption, like human males do (La Vignera, Condorelli, Balercia, Vicari, & Calogero, 2013; Salas-Huetos, Bulló, & Salas-Salvadó, 2017). GB47217 is a calcitonin receptor and is involved in temperature preference rhythm regulation and body temperature preference rhythms during active phases in Drosophila melanogaster and mice respectively (Goda et al., 2018). Hypomethylation of the calcitonin receptor is involved in aggression behavior in honey bees (Herb et al., 2018). Our results show that ethanol induces hypermethylation of the calcitonin receptor, thus it is unlikely that ethanol would invoke a state of aggression in honey bees, speculatively rather resulting in more sedate behavior that could possibly have a negative effect on life span. GB43904 is part of the IFT-B complex that binds to tubulin and mediates tubulin transport within the cilium (Duran et al., 2016) (Bhogaraju et al., 2013).

In the one region that ethanol induced hypomethylation, curcumin co-consumption failed to fully protect against this hypomethylation, however we observed a trend of partial protection (Figure 3D). This region is spanning part of the gene body of locus GB52826, a transmembrane protein that seems to be ubiquitously expressed in human tissues and associated with thyroid cancers (Akaishi et al., 2007) (Fagerberg et al., 2014).

In summary, all the loci that were affected by DNA hypermethylation as a result of ethanol consumption showed full or partial protection when curcumin was consumed together with ethanol. Our results indicate few, but nevertheless robust curcumin-ethanol interactions on the level of methylation changes, suggesting a function for curcumin in blocking the effect of ethanol on DNA methylation. In humans, ethanol has been shown to alter DNA methylation patterns in adults that persist long after the substance abuse has stopped (Cobben et al., 2019). These changes can also affect the germ line, suggesting that these changes can be transmitted to the next generation (Govorko, Bekdash, Zhang, & Sarkar, 2012). Further studies are needed to investigate if curcumin can protect against ethanol associated DNA methylation changes during or following ethanol abuse in humans.

Furthermore, we identified region specific DNA methylation changes as a result of ethanol intake. Ethanol specific changes in DNA methylation were fully or partially blocked in honey bees receiving ethanol and curcumin together. Ethanol-affected and curcumin-blocked differentially methylated regions covered genes involved in fertility, temperature regulation and tubulin transport. Our results demonstrate fundamental negative effects of low dose ethanol consumption on lifespan and associated DNA methylation changes and present a proof-of-principle on how longevity and DNA methylation changes can be restored by the bioactive food component curcumin.

Our findings provide a fundament for further studies of curcumin in mice and humans and offer an avenue to explore regarding possible prevention of health issues related to alcohol consumption.

## Conclusions

In summary, our results show that bioactive food components of relevance for humans can modulate lifespan in the honey bee. Our data opens up for the possibility that negative effects of ethanol on lifespan might be facilitated trough changes in DNA methylation, and that these changes are prevented when curcumin is consumed together with ethanol. We present a proof-of-principle on how longevity and DNA methylation changes can be restored by the bioactive food component curcumin. Furthermore, we extend the honey bee as model system for the study of DNA methylation changes in relation to alcoholism by introducing an intervention with curcumin. Our findings provide a fundament for further studies of curcumin in mice and humans and offer an avenue to explore regarding possible prevention of health issues related to alcohol consumption. It would be of uttermost value to the overall health and healthspan of numerous human societies to identify a compound that can significantly reduce the negative effects of alcohol consumption on human health.

## Acknowledgements

The authors thank Dr Brian Herb for sharing his R scripts, professor Bing Ren for carrying out sequencing at his laboratory, and Professor Peter Aleström for his assistance in the manuscript preparation.

## Author contributions

DM, EMKR, JAD and GVA designed the study. KLS, IKP, CK and EMKR carried out the animal experiments and KLS and IKP performed the ELISA. EMKR constructed the sequencing libraries. EMKR analyzed the data and generated the figures with support from JAD, DM and GVA. EMKR wrote the manuscript with assistance from GVA, DM and JAD. All authors read and approved the manuscript before publication.

