## Supplemental figures 1-11 for "Screening bioactive food compounds in honey bees suggests curcumin blocks alcohol-induced damage to longevity and DNA methylation"

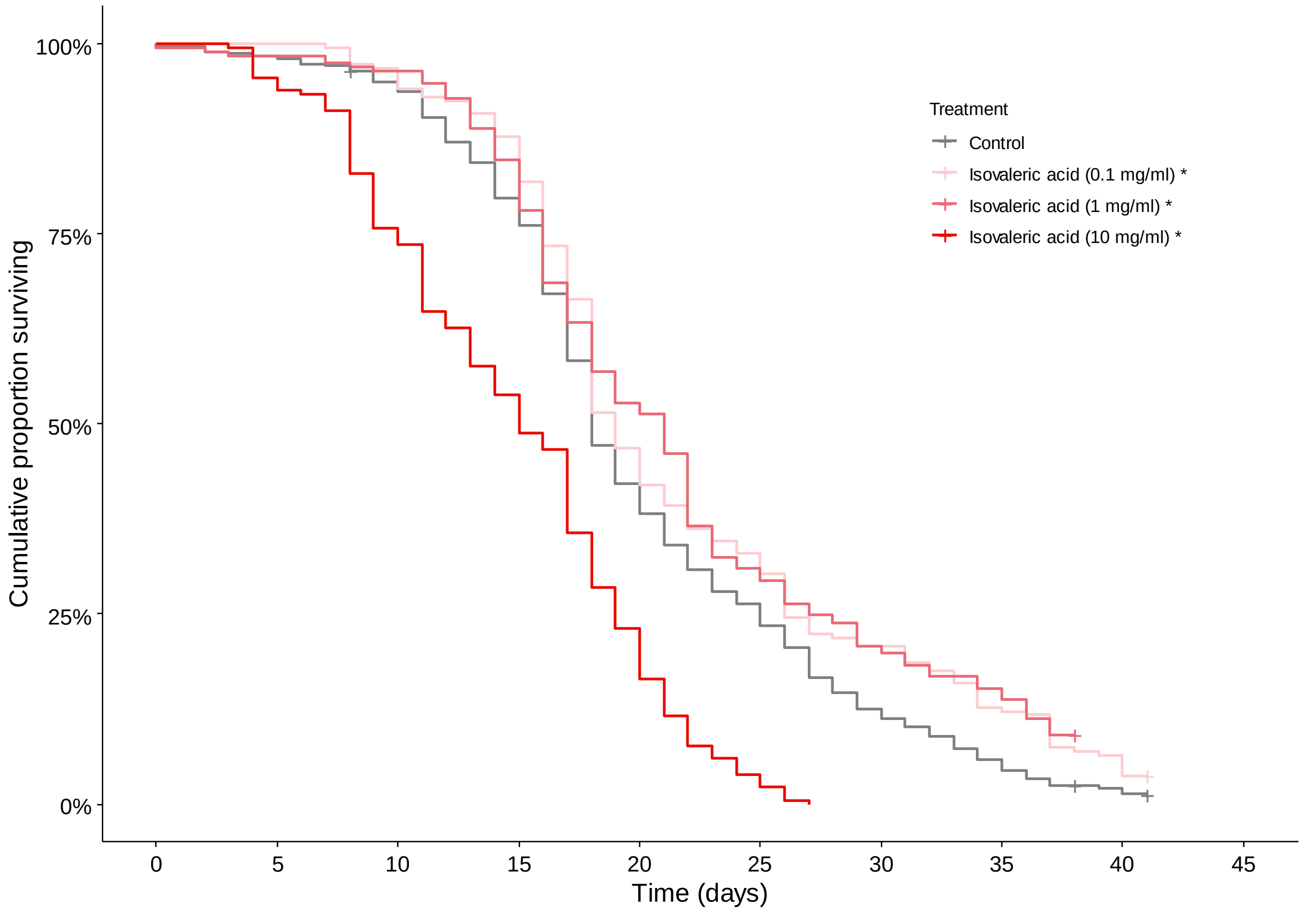


Supplemental figure 1: Cumulative proportion of surviving honey bees exposed to isovaleric acid in feed at 10 mg/ml, 1 mg/ml and 0.1 mg/ml. Pluss signs indicate censored honey bees. Asterisks in legends indicates significant effects on survival compared to control ($P < 0.05$, Pair-wise Log-Rank test corrected using FDR).

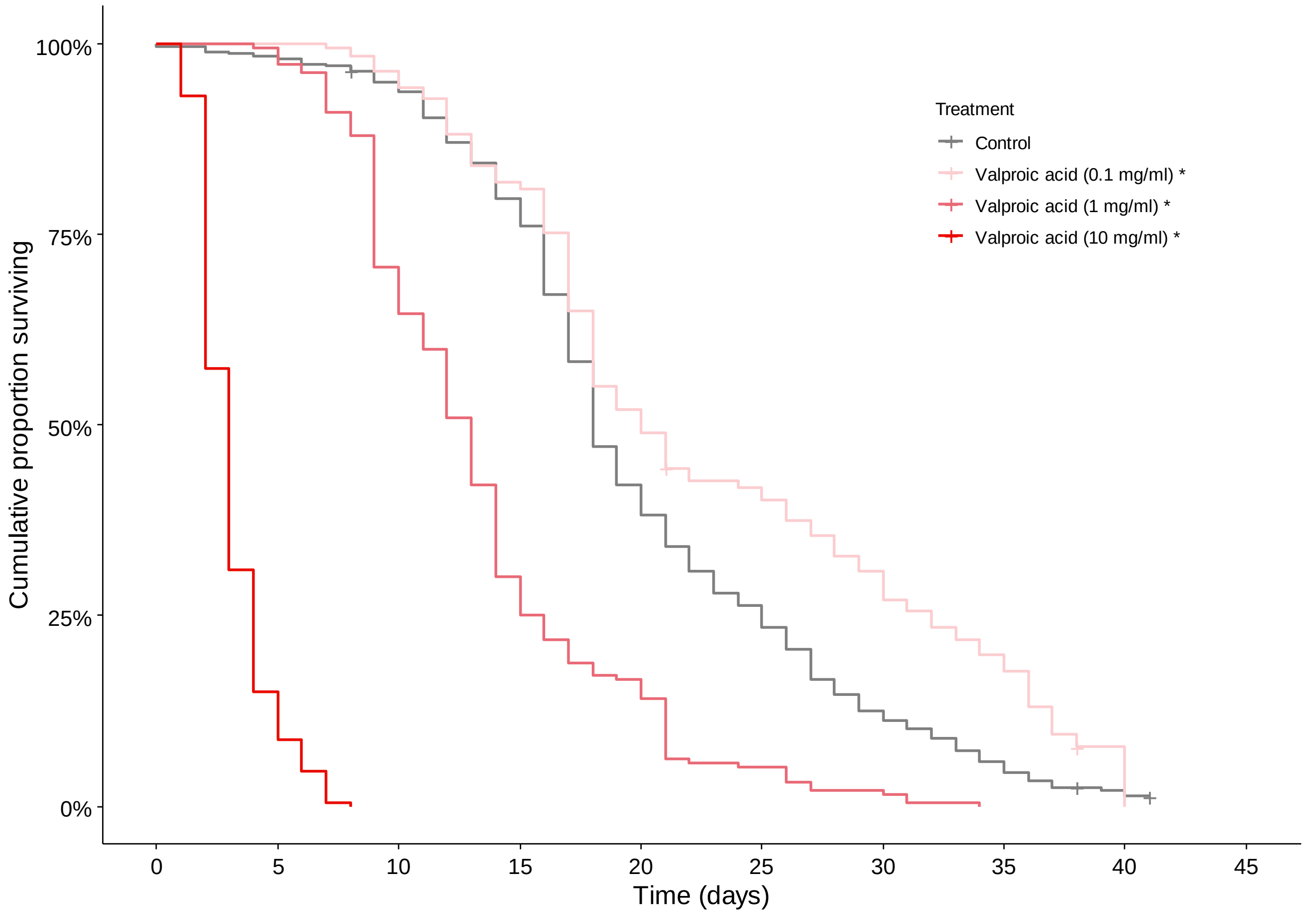


Supplemental figure 2: Cumulative proportion of surviving honey bees exposed to Valproic acid in feed at 10 mg/ml, 1 mg/ml and 0.1 mg/ml. Pluss signs indicate censored honey bees. Asterisks in legends indicates significant effects on survival compared to control (P < 0.05, Pair-wise Log-Rank test corrected using FDR).

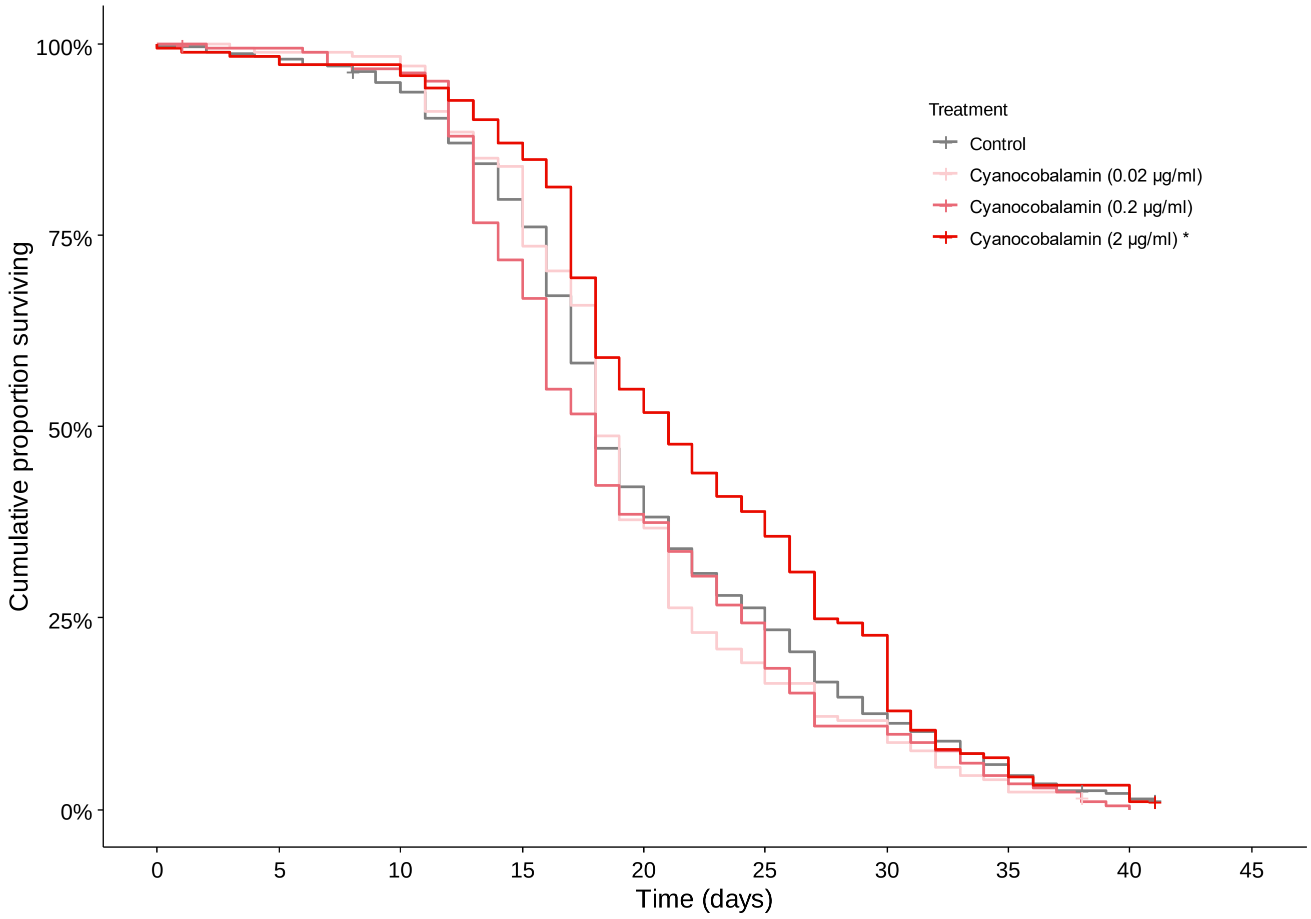


Supplemental figure 3: Cumulative proportion of surviving honey bees exposed to cyanocobalamin in feed at 2 µg/ml, 0.2 µg/ml and 0.02 µg/ml. Pluss signs indicate censored honey bees. Asterisks in legends indicates significant effects on survival compared to control (P < 0.05, Pair-wise Log-Rank test corrected using FDR).

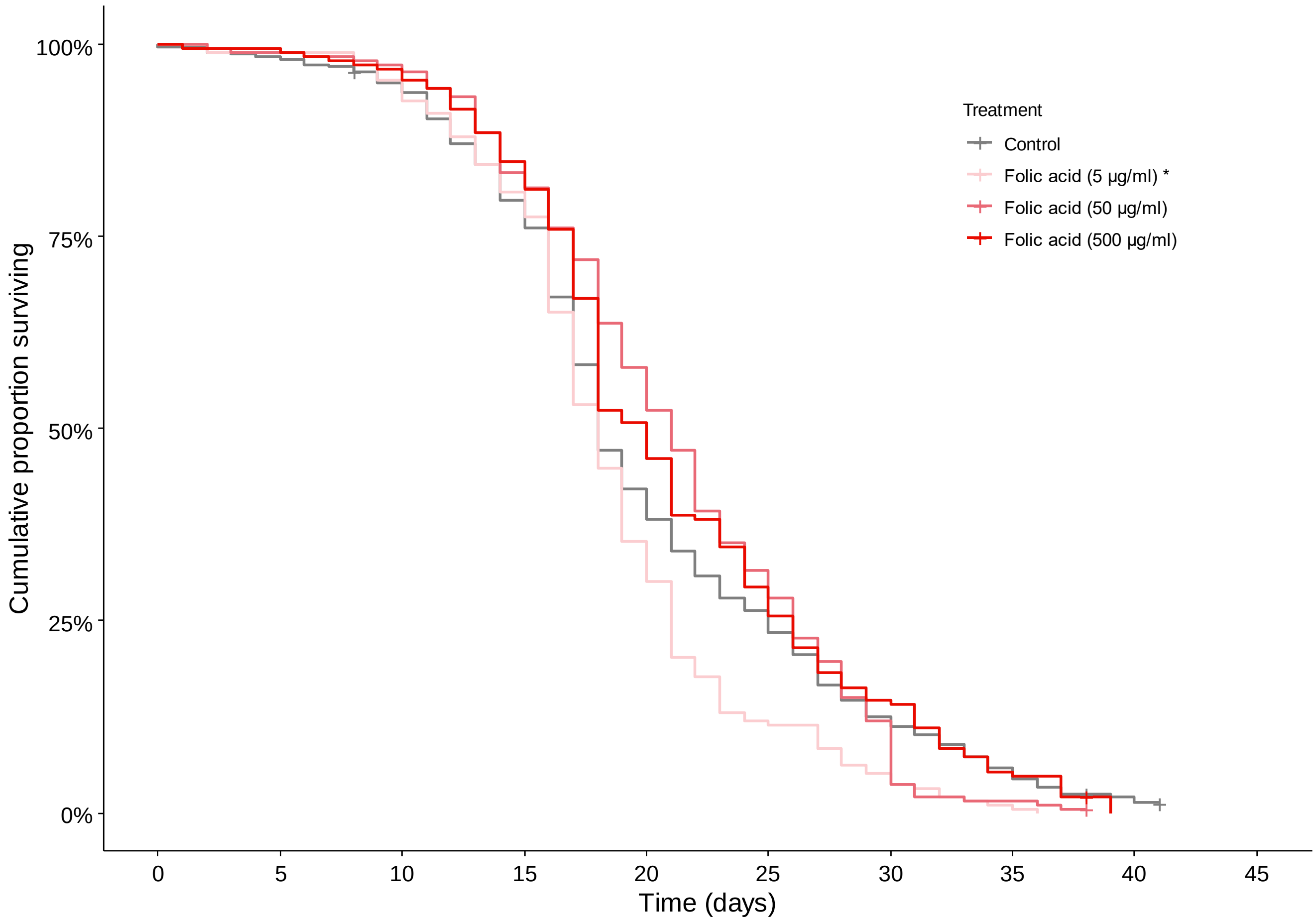


Supplemental figure 4: Cumulative proportion of surviving honey bees exposed to folic acid in feed at 500 µg/ml, 50 µg/ml and 5 µg/ml. Pluss signs indicate censored honey bees. Asterisks in legends indicates significant effects on survival compared to control ($P < 0.05$, Pair-wise Log-Rank test corrected using FDR).

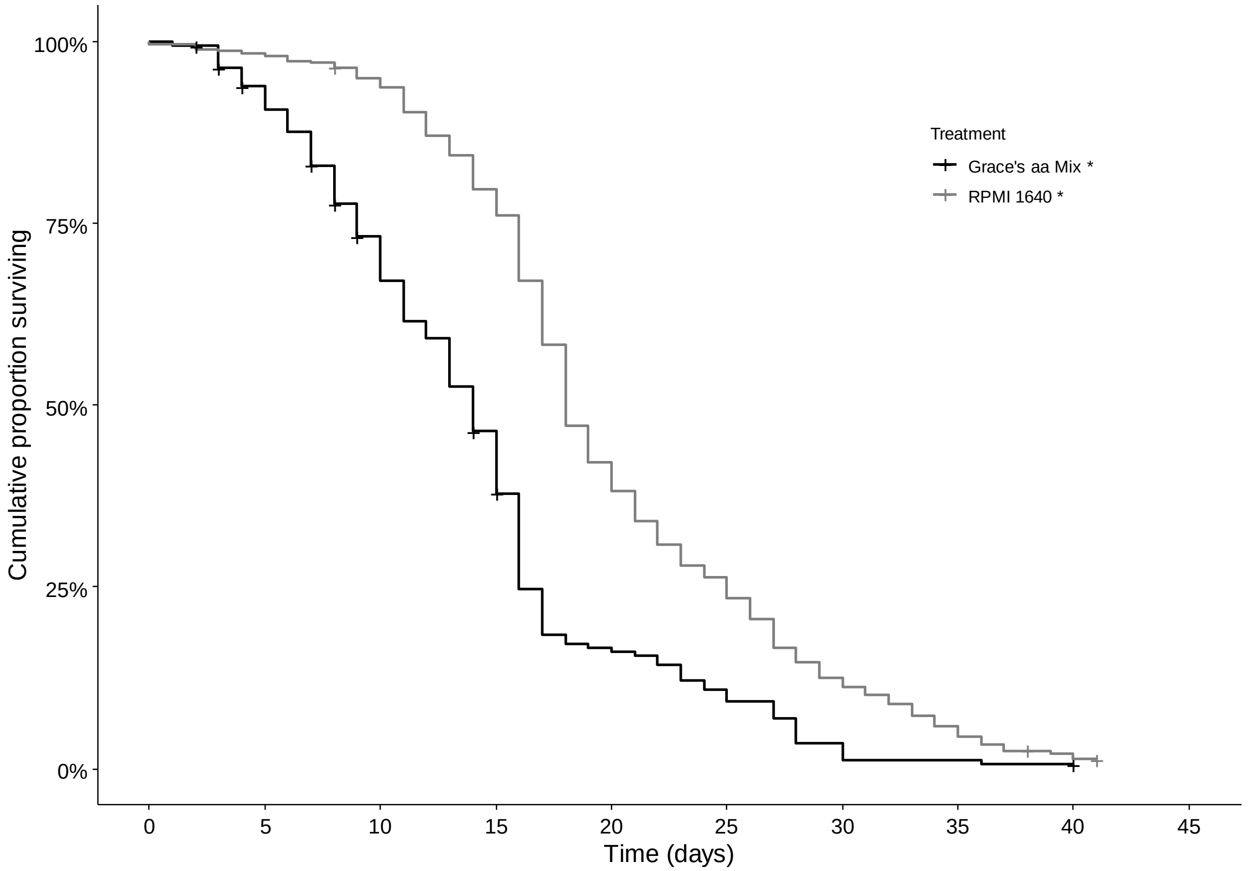


Supplemental figure 5: Cumulative proportion of surviving honey bees exposed to different sources of amino acids. Pluss signs indicate censored honey bees. Asterisks in legends indicates significant effects on survival (Kaplan Meier, $p < 0.0001$).

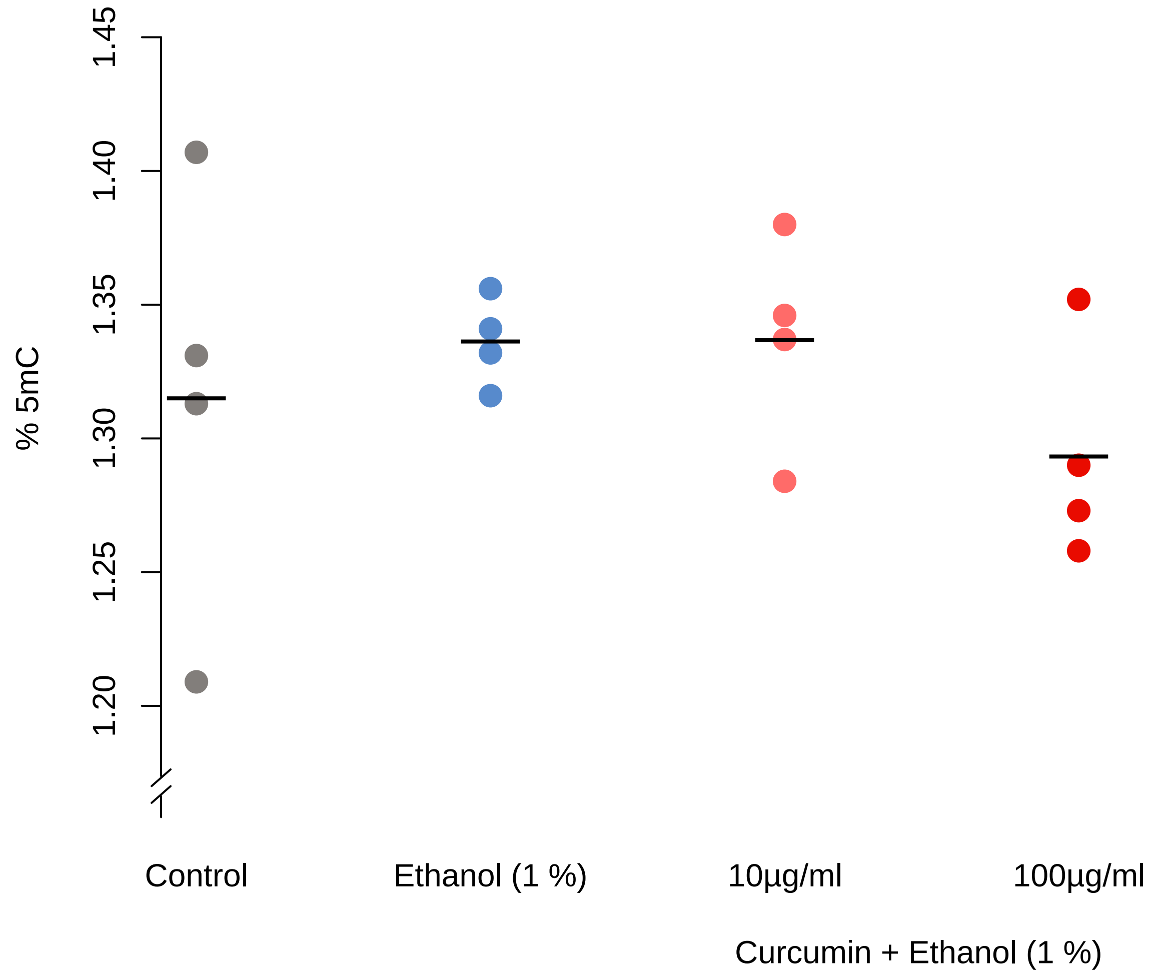


Supplemental figure 6: ELISA methylation percentages. Black lines indicate means. (One−tailed Mann−Whitney test; $p>0.05$)

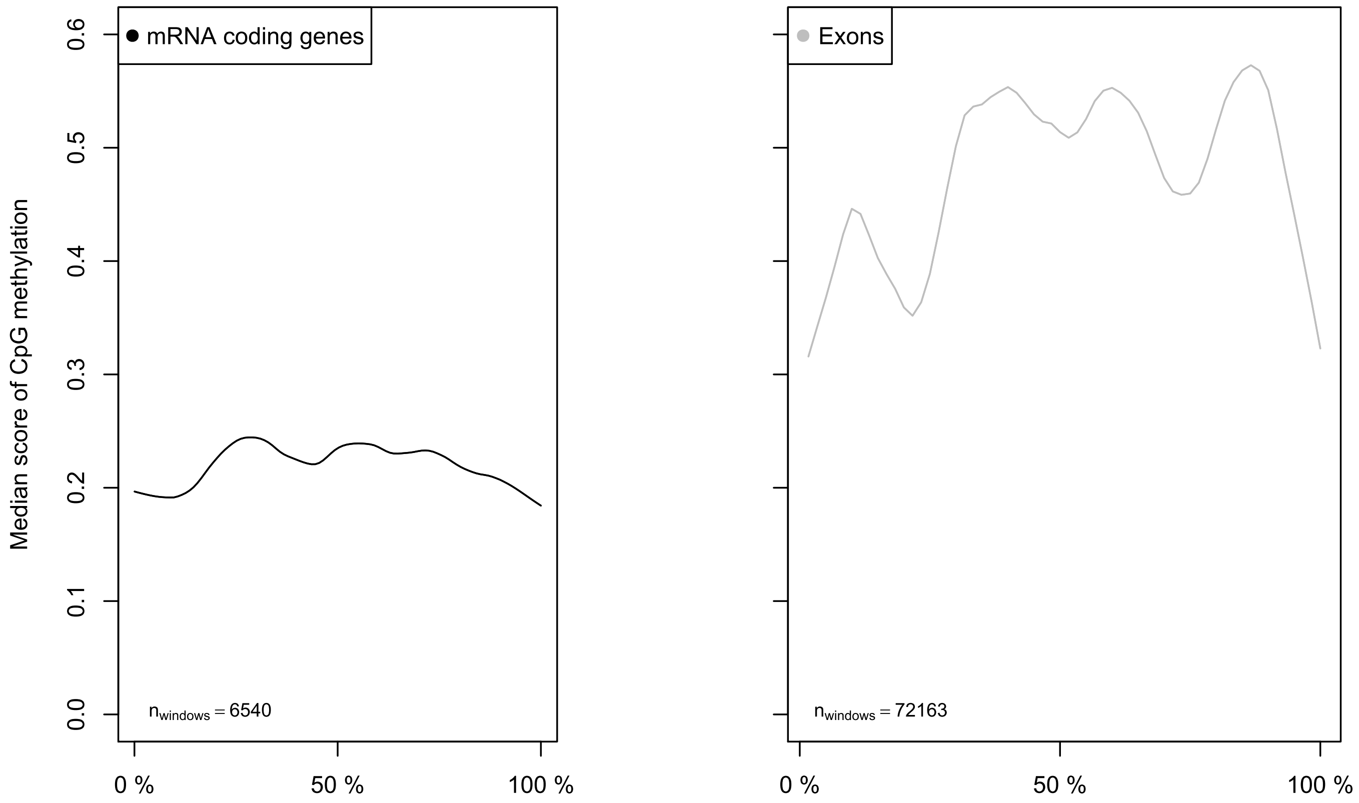


Supplemental figure 7: Median CpG methylation score over mRNA coding genes (left panel) and exons (right panel). mRNA coding genes and exons were divided into a minimum of 3,000 and 60 bins respectively. Scores are re-scaled in the range from 0 to 1 and lines are smoothed using Locally Weighted Scatterplot Smoothing (LOWESS).

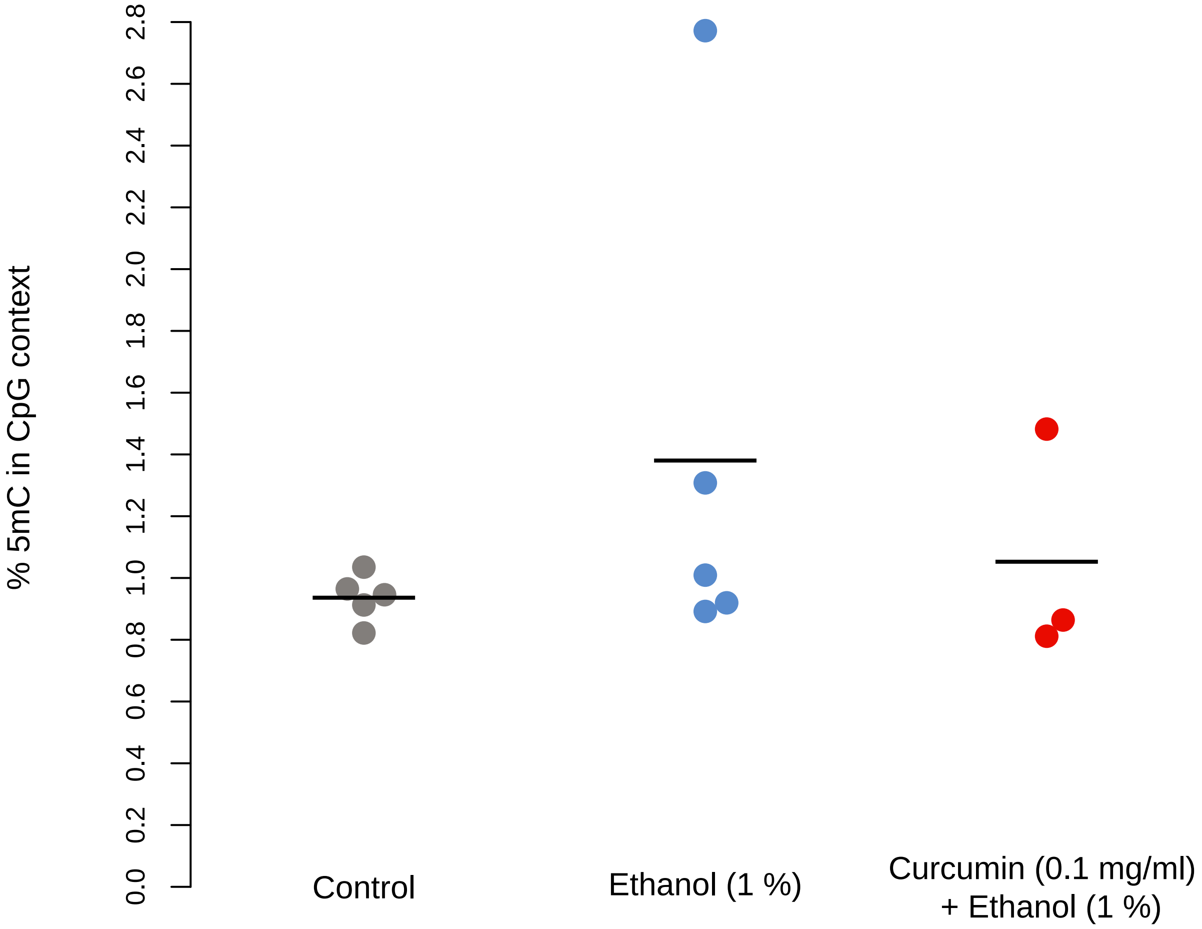


Supplemental figure 8: WGBS CpG methylation percentages. Black lines indicate means.
(One−tailed Mann−Whitney tests, $p > 0.05$)

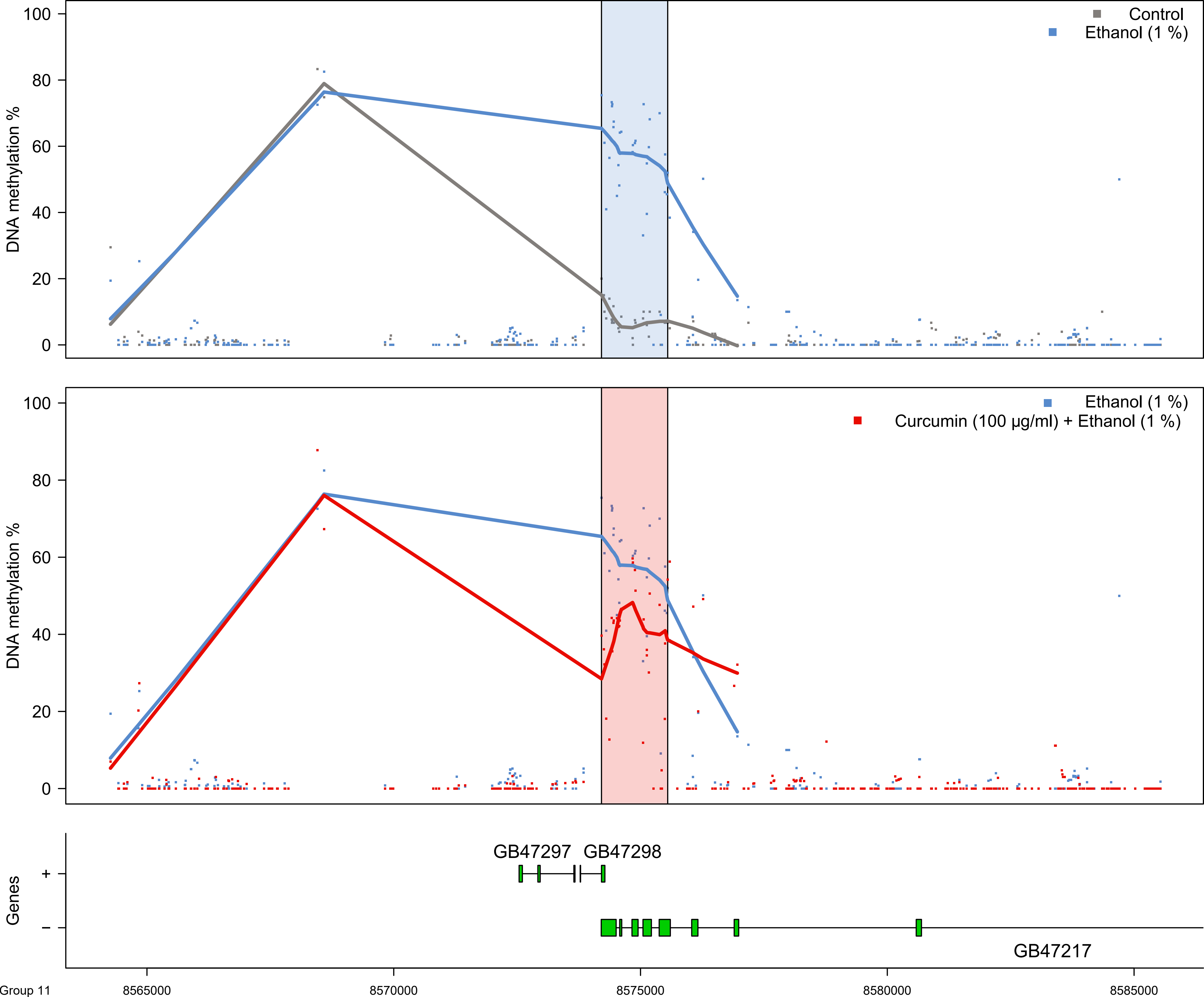

Control
Ethanol (1 %)
DNA methylation %
100
80
60
40
20
0
Ethanol (1 %)
Curcumin (100 µg/ml) + Ethanol (1 %)
DNA methylation %
100
80
60
40
20
0
GB47297
GB47298
Genes
+
–
GB47217
Group 11
8565000
8570000
8575000
8580000
8585000

Supplemental figure 9: Genomic regions of the DMR at linkage group 11 where the top panel depicts the situation in control (grey) versus ethanol (blue) and the blue shaded area corresponds to the significant DMR in the middle panel. 10 kbp upstream and downstream (if no gaps exist in the Amel 4.5 genome build) are plotted for reference (white background). The middle panel depicts the comparison between ethanol (blue) and the curcumin and ethanol (red) fed honey bees where the significant DMR is in the pink shaded area. The lower panel depicts the gene structures by exons (boxes) by strand according to linkage groups.

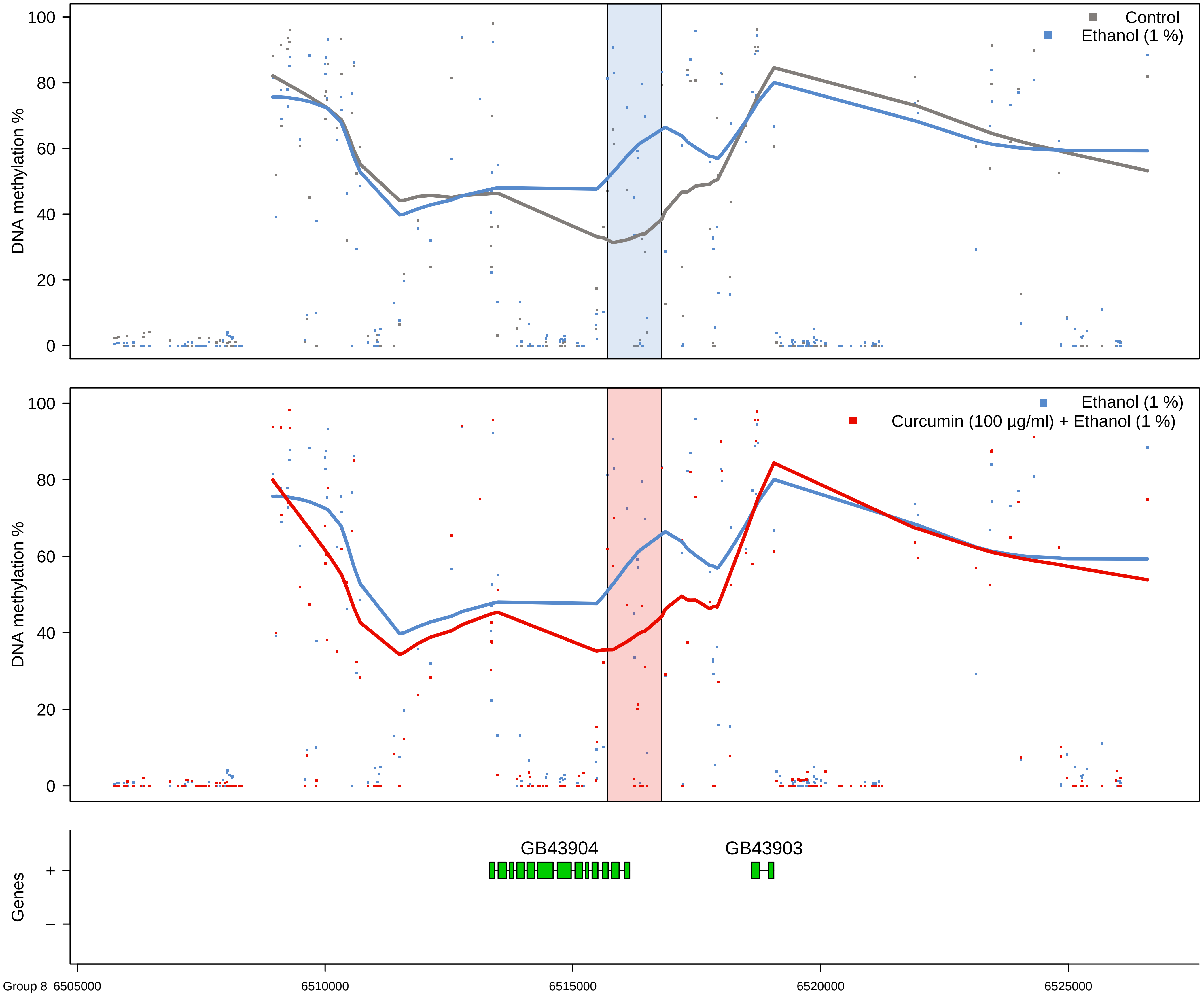

Control
Ethanol (1 %)
DNA methylation %
Curcumin (100 µg/ml) + Ethanol (1 %)
GB43904
GB43903
Genes
+
−
Group 8 6505000
6510000
6515000
6520000
6525000

Supplemental figure 10: Genomic regions of the DMR at linkage group 8 where the top panel depicts the situation in control (grey) versus ethanol (blue) and the blue shaded area corresponds to the significant DMR in the middle panel. 10 kbp upstream and downstream (if no gaps exist in the Amel 4.5 genome build) are plotted for reference (white background). The middle panel depicts the comparison between ethanol (blue) and the curcumin and ethanol (red) fed honey bees where the significant DMR is in the pink shaded area. The lower panel depicts the gene structures by exons (boxes) by strand according to linkage groups.

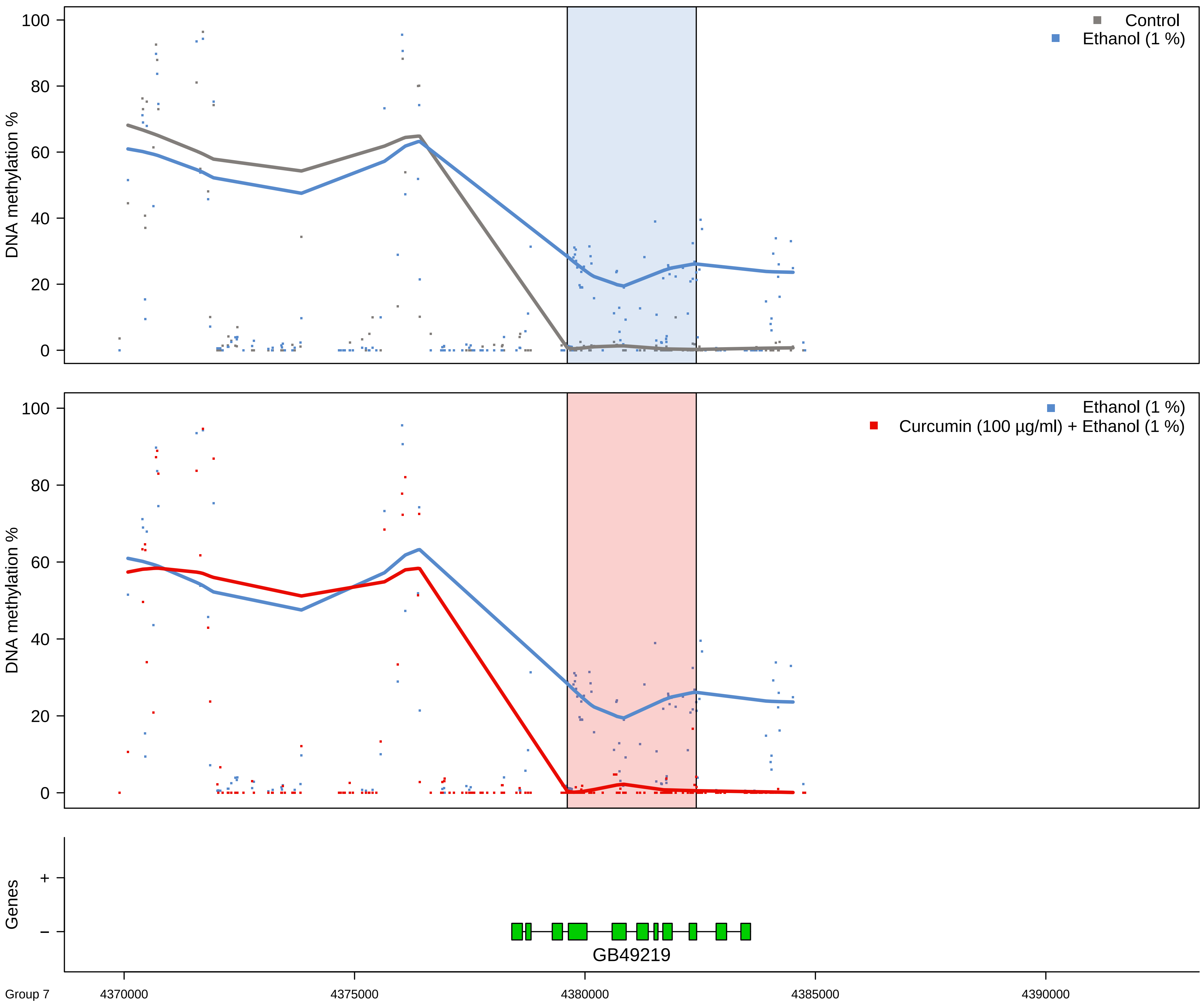

Control
Ethanol (1 %)
DNA methylation %
0
20
40
60
80
100
Ethanol (1 %)
Curcumin (100 µg/ml) + Ethanol (1 %)
DNA methylation %
Genes
+
−
GB49219
Group 7
4370000
4375000
4380000
4385000
4390000

Supplemental figure 11: Genomic regions of the DMR at linkage group 7 where the top panel depicts the situation in control (grey) versus ethanol (blue) and the blue shaded area corresponds to the significant DMR in the middle panel. 10 kbp upstream and downstream (if no gaps exist in the Amel 4.5 genome build) are plotted for reference (white background). The middle panel depicts the comparison between ethanol (blue) and the curcumin and ethanol (red) fed honey bees where the significant DMR is in the pink shaded area. The lower panel depicts the gene structures by exons (boxes) by strand according to linkage groups.
