## Supplemental Table 1 for "Screening bioactive food compounds in honey bees suggests curcumin blocks alcohol-induced damage to longevity and DNA methylation"

| **Amino acid source** | **Compounds** | **Substance solvent** |
| --- | --- | --- |
| Grace’s amino acid mix | Sodium butyrate 0.01 mg/ml | dH_2_O |
| **Grace’s amino acid mix** | **Sodium butyrate 0.1 mg/ml** | **dH_2_O** |
| **Grace’s amino acid mix** | **Sodium butyrate 1 mg/ml** | **dH_2_O** |
| Grace’s amino acid mix | None | dH_2_O |
| **Grace’s amino acid mix** | **Ethanol 1 % (v/v)** | **dH_2_O** |
| Grace’s amino acid mix | Curcumin 1 µg/ml | Ethanol 1 % (v/v) |
| **Grace’s amino acid mix** | **Curcumin 10 µg/ml** | **Ethanol 1 % (v/v)** |
| **Grace’s amino acid mix** | **Curcumin 100 µg/ml** | **Ethanol 1 % (v/v)** |
| **RPMI 1640** | **Cyanocobalamin 0.02 µg/ml** | **dH_2_O** |
| **RPMI 1640** | **Cyanocobalamin 0.2 µg/ml** | **dH_2_O** |
| RPMI 1640 | Cyanocobalamin 2 µg/ml | dH_2_O |
| **RPMI 1640** | **Folic acid 5 µg/ml** | **dH_2_O** |
| RPMI 1640 | Folic acid 50 µg/ml | dH_2_O |
| **RPMI 1640** | **Folic acid 500 µg/ml** | **dH_2_O** |
| **RPMI 1640** | **Valproic acid 0.1 mg/ml** | **dH_2_O** |
| **RPMI 1640** | **Valproic acid 1 mg/ml** | **dH_2_O** |
| RPMI 1640 | Valproic acid 10 mg/ml | dH_2_O |
| RPMI 1640 | Isovaleric acid 0.1 mg/ml | dH_2_O |
| **RPMI 1640** | **Isovaleric acid 1 mg/ml** | **dH_2_O** |
| **RPMI 1640** | **Isovaleric acid 10 mg/ml** | **dH_2_O** |
| **RPMI 1640** | **None** | **dH_2_O** |

Supplementary table 1: Detailed makeup of honey bee diets. Compound concentrations in bold were used for ELISA experiments.
